# Efficient querying of genomic reference databases with *gget*

**DOI:** 10.1101/2022.05.17.492392

**Authors:** Laura Luebbert, Lior Pachter

## Abstract

**Motivation:** A recurring challenge in interpreting genomic data is the assessment of results in the context of existing reference databases. Currently, there is no tool implementing automated, easy programmatic access to curated reference information stored in a diverse collection of large, public genomic databases.

**Results:** *gget* is a free and open-source command-line tool and Python package that enables efficient querying of genomic reference databases, such as Ensembl. *gget* consists of a collection of separate but interoperable modules, each designed to facilitate one type of database querying required for genomic data analysis in a single line of code.

**Availability:** The manual and source code are available at https://github.com/pachterlab/gget.

## Introduction

The increasingly common use of genomic methods, such as single-cell RNA-seq, to provide transcriptomic characterization of cells is dependent on quick and easy access to reference information stored in large genomic databases such as Ensembl, NCBI, and UniProt (Cunningham et al., 2022; NCBI Resource Coordinators, 2013; UniProt Consortium, 2021). Although integrated information retrieval systems date back to the 1990s (Etzold et al., 1996; Zdobnov et al., 2002), a majority of researchers currently access genomic reference databases to annotate and functionally characterize putative marker genes through web access (Stalker et al., 2004; Birney et al., 2004). This process is time-consuming and error-prone, as it requires manually copying and pasting data, such as gene IDs.

To facilitate and automate functional annotation for genomic data analyses, we developed *gget:* a free and open-source software package that rapidly queries information stored in several large, public databases directly from a command line or Python environment. *gget* consists of a collection of tools designed to perform the database querying required for genomic data analysis in a single line of code. In addition to providing access to genomic databases, *gget* can also leverage sequence analysis tools, such as BLAST (Altschul et al., 1990, 1997), thus simplifying complex annotation workflows.

While there are some web-based Application Programming Interface (API) data mining systems, such as BioMart (Durinck et al., 2005; Kasprzyk et al., 2004), we identified several limitations in such tools, including limits to query types and to utilizing databases in tandem. Moreover, large-scale genomic data analyses, such as single-cell RNA-seq data analysis, are better served by command line or packaged APIs that can fetch data directly into programming environments.

The *gget* modules combine MySQL (Oracle Corporation, 1995), API, and web data extraction queries to rapidly and reliably request comprehensive information from different databases (Figure 1). This approach allows *gget* to perform tasks unsupported by existing tools built around standard API queries (de Ruiter, 2016). For instance, searching for genes and transcripts using free-form search terms. Each *gget* tool requires minimal arguments, provides clear output, and operates from both the command line and Python environments, such as JupyterLab, maximizing ease of use and accommodating novice programmers.

**Figure 1.**
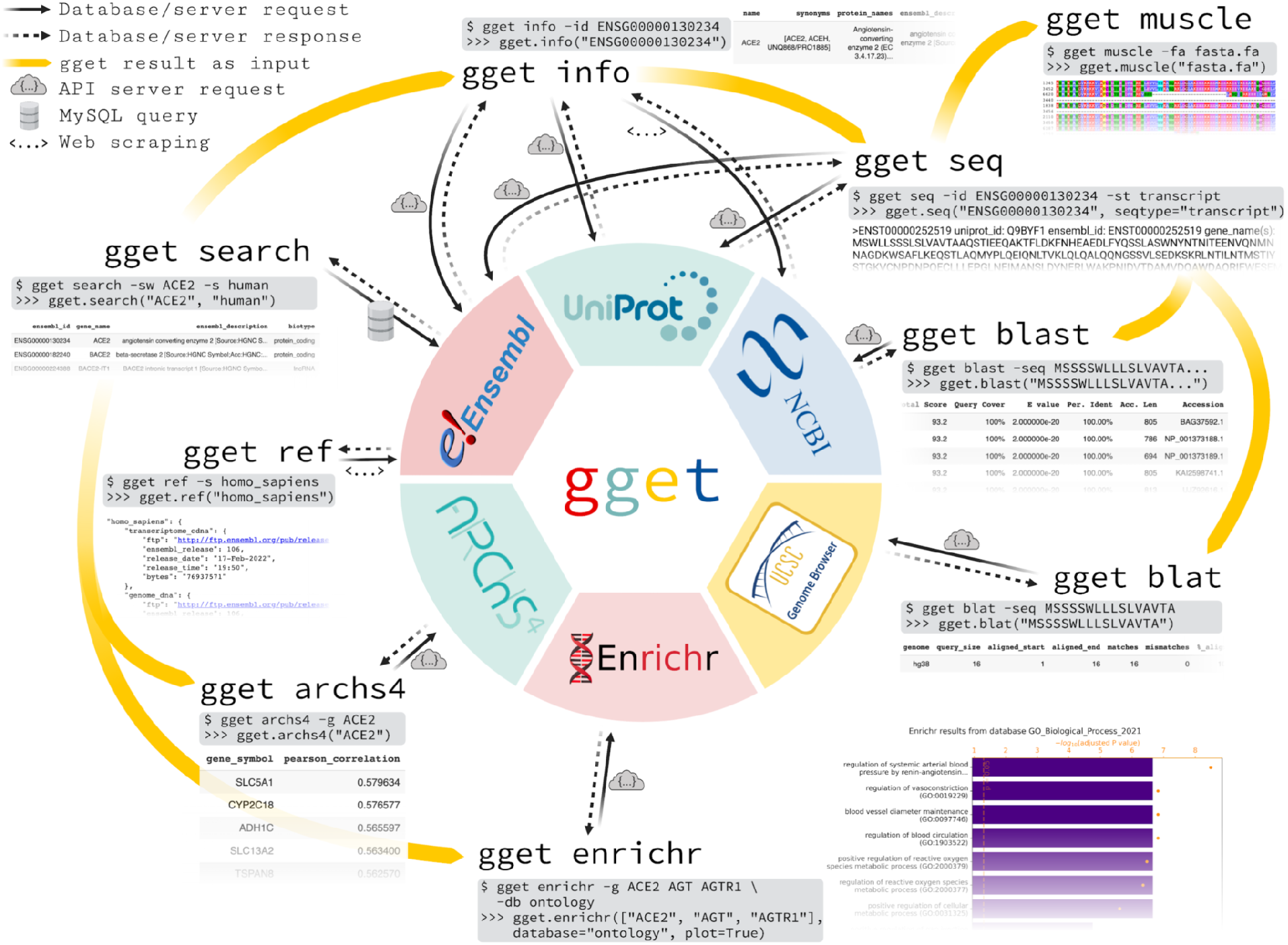
Overview of the nine *gget* tools and the public databases they access. One simple command line ($) example and its Python (>>>) equivalent are shown for each tool with the corresponding output.

## Description

*gget* consists of nine tools:

- *gget ref:* Fetch File Transfer Protocols (FTPs) and metadata for reference genomes or annotations from Ensembl by species.
- *gget search:* Fetch genes or transcripts from Ensembl using free-form search terms.
- *gget info:* Fetch extensive gene or transcript metadata from Ensembl, UniProt, and NCBI by Ensembl ID.
- *gget seq:* Fetch nucleotide or amino acid sequences of genes or transcripts from Ensembl or UniProt by Ensembl ID.
- *gget blast:* BLAST (Altschul et al., 1990, 1997) a nucleotide or amino acid sequence to any BLAST database.
- *gget blat:* Find the genomic location of a nucleotide or amino acid sequence using BLAT (James Kent, 2002).
- *gget muscle:* Align multiple nucleotide or amino acid sequences to each other using the Muscle5 algorithm (Edgar, 2021).
- *gget enrichr:* Perform an enrichment analysis on a list of genes using Enrichr (Chen et al., 2013; Xie et al., 2021; Kuleshov et al., 2016) and an extensive collection of gene set libraries, including KEGG (Kanehisa and Goto, 2000; Kanehisa, 2019; Kanehisa et al., 2021) and Gene Ontology (Ashburner et al., 2000; Gene Ontology Consortium, 2021).
- *gget archs4:* Find the most correlated genes to a gene of interest or find the gene’s tissue expression atlas using ARCHS4 (Lachmann et al., 2018).

Each *gget* tool accesses data stored in one or several public databases, as depicted in Figure 1. *gget* fetches the requested data in real-time, guaranteeing that each query will return the latest information. One exception is *gget muscle,* which locally compiles the Muscle5 algorithm (Edgar, 2021) and therefore does not require an internet connection.

*gget info* combines information from Ensembl, NCBI, and UniProt (Cunningham et al., 2022; NCBI Resource Coordinators, 2013; UniProt Consortium, 2021) to provide the user with a comprehensive executive summary of the available information about a gene or transcript. This also enables users to assert whether data from different sources are consistent.

By accessing the NCBI server (NCBI Resource Coordinators, 2013) through HTTP requests, *gget blast* does not require the download of a reference BLAST database, as is the case with existing BLAST tools (Buchfink et al., 2021; Camacho et al., 2009). The whole self-contained *gget* package is approximately 3 MB after installation.

The package dependencies were carefully chosen and kept to a minimum. *gget* depends on the HTML parser *beautifulsoup4* (Richardson, 2022), the Python MySQL-connector (Oracle, 2022), and the HTTP library *requests* (Reitz, 2022). All of these are well-established packages for server interaction in Python. *gget* has been tested on Linux/Unix, Mac OS (Darwin), and Windows.

## Usage and documentation

*gget* can be installed from the command line by running ‘pip install gget’. Figure 1 depicts one use case for each *gget* tool with the corresponding output.

Each *gget* tool features an extensive manual available as function documentation in a Python environment or as standard output using the help flag [-h] in the command line. The complete manual with examples can be viewed in the *gget* repository, available at https://github.com/pachterlab/gget. A separate *gget examples* repository is accessible at https://github.com/pachterlab/gget_examples and includes exemplary workflows immediately executable in Google Colaboratory (Bisong, 2019).

## Discussion

Our open-source Python and command-line program *gget* enables efficient and easy programmatic access to information stored in a diverse collection of large, public genomic reference databases. *gget* works alongside existing tools that fetch user-generated sequencing data (Gálvez-Merchán et al., 2022) to replace ineffective, error-prone manual web access during genomic data analysis. While the *gget* modules were motivated by experience with tedious single-cell RNA-seq data analysis tasks (Supplementary Figure 1), we anticipate their utility for a wide range of bioinformatics tasks.

## Acknowledgments

We thank Kyung Hoi (Joseph) Min for advice on the command-line interface, Matteo Guareschi for advice on Windows operability, and A. Sina Booeshaghi, Kristján Eldjárn Hjörleifsson, and Ángel Gálvez-Merchán for insightful discussions about *gget*. Illustrations in Figure 1 and Supplementary Figure 1 were created with BioRender.com. Thanks to the wonderful staff at Dash Coffee Bar in Pasadena, who occasionally gave LL free banana bread to sustain this work.

