## Supplementary Information for "Efficient querying of genomic reference databases with *gget*"

### Supplementary Information:
### Efficient querying of genomic reference databases with *gget*


Laura Luebbert[1] and Lior Pachter[1,2,*]

1 Division of Biology and Biological Engineering, California Institute of Technology, Pasadena, California
2 Department of Computing and Mathematical Sciences, California Institute of Technology, Pasadena, California


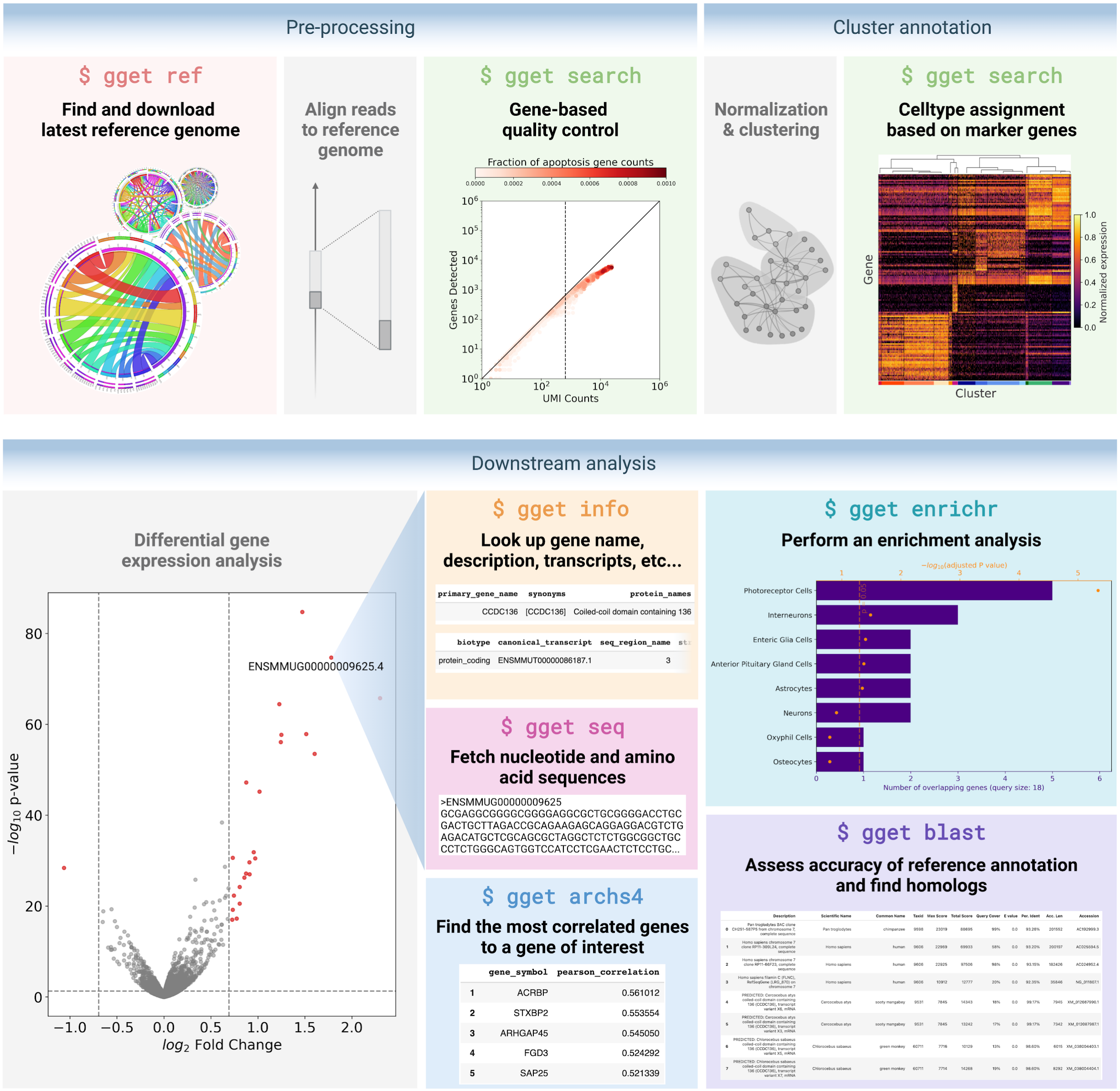


**Supplementary Figure 1** *gget* performs the database querying underlying a standard single-cell RNA-seq data analysis workflow. The workflow and all of the figures are reproducible starting with raw reads using Google Colaboratory notebooks that can be run for free and are accessible at https://github.com/pachterlab/gget_examples/tree/main/scRNAseq_workflow.
